# Distinct neural temporal architectures encode rapid social expressions and sustained internal mood states

**DOI:** 10.64898/2025.12.20.692681

**Authors:** Bina Kakusa, Jay R. Gopal, Lillian Forman, James W. Hartford, Tristan Brennan, Sofia Pantis, Amit R. Persad, Christopher C. Cline, Vivek P. Buch, Josef Parvizi, Yuhao Huang, Corey J. Keller

## Abstract

Affective processing operates across multiple temporal scales, from rapid social signaling through facial expressions to sustained internal mood states, yet the neural computational principles governing these different timescales remain unclear. Understanding how the brain implements distinct temporal architectures for momentary versus persistent affective phenomena is important to comprehending emotional processing and developing objective biomarkers for psychiatric conditions. Here, we introduced a multimodal approach combining automated facial expression monitoring and continuous intracranial electroencephalography in 2,037 electrode contacts across 16 epilepsy patients, over multiple days. Of these, 15 and 12 patients met criteria for facial expression and for mood analysis, respectively. Among patients meeting criteria, we captured 1,396 naturalistic smiles, and 3,746 neutral expressions – separated by at least 10 seconds, alongside 336 periodic mood assessments. This paradigm revealed distinct behavioral and neural computational architectures. Aperiodic neural activity in the lateral temporal cortex (79.5% accuracy) encoded facial expressions with high cross-participant generalizability. Mood states, however, showed different encoding patterns. Facial expressions provided no consistent mood indicators across participants. Critically, low-gamma power dynamics in limbic regions encoded mood states in only a subset of individuals (5 of 12 participants) with expression-mood behavioral correlations, suggesting a distinct encoding phenotype. Cross-domain analysis confirmed computational independence: neural features optimized for facial expression decoding failed to predict sustained mood states, and vice versa. These findings suggest that multiple neural mechanisms may influence underlying affective processing, with variations in their contributions between individuals. The results provide a framework for understanding individual differences in neural mood representation and establish methodological approaches for objective measurement of naturalistic affective behaviors.

## Introduction

Affective processing in the human brain operates across different temporal scales, from rapid social signaling through facial expressions to sustained internal mood states. The distinct neural computational architecture governing these temporal domains remains poorly understood. Current neuroscience frameworks often assume shared substrates between momentary expressions and persistent affective states, but whether the brain implements separate temporal architectures for these phenomena represents a critical gap in our understanding of emotional processing^1^. While facial expressions provide observable windows into rapid social communication occurring over seconds^2^, internal mood states reflect sustained neural configurations operating over minutes to hours^3,4^. The temporal dissociation between these systems has profound implications for developing objective neural biomarkers for psychiatric disorders^5,6^ and understanding individual differences in affective representation, as current neuropsychiatric practice relies heavily on subjective self-reports that conflate these distinct temporal processes^7^. Determining whether rapid social expressions and sustained mood states involve distinct or overlapping neural processes is essential for advancing precision approaches to mental health assessment and treatment^4^.

Facial expressions offer a continuous and observable behavioral window into emotional processes, and have been studied extensively using both facial action coding systems^8^ and computer vision approaches^9^. Clinical studies have linked facial expressions to psychiatric disorders including depression^8^ and anxiety^10,11^, and representations of internal experiences such as pain^12^. Facial expression processing involves rapid, distributed neural computations across temporal-limbic networks^13^. Neuroimaging studies show that distinct facial expressions engage dynamic functional connectivity across distributed networks, including prefrontal, insular, superior temporal gyrus, middle temporal gyrus, and fusiform cortex, alongside premotor/orofacial sensorimotor regions, reflecting the coordinated interplay of sensory, emotion, and motor systems^14^. Disruptions in facial expression processing are hallmarks of psychiatric disorders, with altered neural responses to emotional faces observed in depression and anxiety^15^. The ability to accurately decode facial expressions from neural activity could therefore provide objective biomarkers for psychiatric conditions and treatment response. Further, these networks demonstrate remarkable consistency across individuals, suggesting evolutionarily conserved mechanisms for social emotional communication^16–18^.

In contrast, mood states emerge from sustained, distributed configurations of limbic-prefrontal networks operating over much longer timescales^3,4^. Converging evidence from resting-state functional MRI (fMRI), task-based fMRI, and lesion studies of mood disorders implicates distributed limbic-prefrontal networks, including the amygdala, hippocampus, ventromedial prefrontal cortex, and anterior cingulate^3,19–21^. Available literature suggests that the distributed limbic-prefrontal architecture, implicated in mood processing, is also recruited during emotion processing when viewing emotional faces^22^, when mimicking viewed facial expressions^23^, and when expressed emotions are congruent with the viewed emotional face^24^. Further, structural and functional abnormalities of these networks in mood disorders such as major depression, alters activation patterns during emotional face recognition when compared to healthy controls^25^. Intracranial stimulation of nodes within these shared networks, such as the cingulate cortex, evokes distinct internal affective states (e.g., mirth, anxiety, agitation) along with coordinated, goal-directed emotional and motor behaviors (e.g., laughter, smiling, crying, grimacing)^26,27^. Focal limbic-prefrontal and fronto–striatal disruptions, such as in Parkinson’s, are associated with disturbances in emotional behavior (flattened affect, pathological laughter/crying, impaired voluntary emotional facial expression) as well as disturbances in mood (apathy, sustained emotional lability and disinhibition)^28–30^.

Unfortunately, there is a lack of robust neurophysiologic studies which simultaneously explore emotional expression and mood encoding. Facial expressions of smiles are among the most robust behavioral readouts of positive affect^16–18^, yet their relationship to ongoing mood is complex: individuals may smile despite dysphoria, or fail to express overt affect despite euthymia^2^. Moreover, facial expressions operate on rapid timescales of seconds, while mood states persist over minutes to hours, suggesting potentially distinct temporal architectures for these phenomena. Scalp electroencephalography (EEG) and magnetoencephalography (MEG) associate mood and behavioral expression with band-limited power changes, particularly in the gamma and alpha–beta ranges^31,32^. However, noninvasive approaches are unable to resolve the rapid, spatial-specific dynamics neural components that may be critical for naturalistic expression and rarely capture spontaneous facial behavior and mood concurrently. Thus, intracranial EEG (iEEG), largely among epilepsy patients, has become an increasingly powerful tool in clinical populations^33,34^. Prior iEEG studies have characterized affective processing primarily using controlled, evoked paradigms with static images or instructed expressions rather than spontaneous, continuous facial behavior^35–38^, leaving a gap in knowledge.

Disentangling the neural encoding of externally expressed affect from internal mood requires measurements that are both temporally precise and anatomically specific during spontaneous, non–task-based behavior. Here, we leveraged continuous iEEG recordings from epilepsy patients to simultaneously decode naturalistic facial expressions and mood states from the same high-resolution neural signals over multiple days. Furthermore, we employ a markerless computer vision approach to continuously quantify affective behaviors and utilize a simple survey to gauge self-reported, internal states in an inpatient clinical setting. Given the intuitive relationship between facial expressions and mood, we hypothesized that their neural encoding mechanisms would converge. Instead, we discovered that smile decoding was achieved with high accuracy through facial expressions and aperiodic activity in the temporal cortex, while mood prediction succeeded only in a subset of individuals through distinct limbic gamma activity, and not with using facial expressions alone. Cross-domain analysis revealed neural dissociation: features optimized for expression decoding failed to predict mood and vice versa. These findings reveal that momentary facial expressions and sustained mood states are likely supported by distinct neural mechanisms operating on distinct temporal scales, challenging assumptions about shared neural substrates and establishing the need for mood-specific rather than expression-derived biomarkers in precision psychiatry.

## Results

### An automated behavioral pipeline validates FAU12 lip corner elevation as generalizable signature for naturalistic smiles

This IRB-approved study involved 16 voluntary, consenting participants (Table S1) who were under multi-day, in-patient, clinical monitoring for their epilepsy using continuous in-room video and intracranial electroencephalography (iEEG) recordings to decode naturalistic facial expressions and self-reported mood states (Figure 1A). This study employed an automated computer vision pipeline specifically designed for continuous behavioral quantification in clinical settings, enabling analysis of naturalistic expressions across multiple days – a scale impractical with manual annotation methods. On average, 8.4 days (standard deviation: 6.3 days) of 24/7 recordings per participant were processed. Using our behavioral quantification pipeline (Figure 1B) that combines real-time face detection, identity verification, and multi-modal feature extraction, followed by manual rating for independent verification of behavioral labels (Figure 2A), a total of 1,396 smiles (range: 39 – 170 per participant) and 3,746 neutral (range 128 – 359 per participant) behaviors were labeled (Table S1). These behaviors occurred in several naturalistic contexts including when interacting with other individuals, using personal devices, watching television, etc. On average across participants, individual facial expressions were separated by 6.03 minutes (SD: 4.9 minutes) with a minimum interval of 10 seconds. One participant was excluded for insufficient number of smiles captured during their inpatient stay to allow analysis, leaving 15 participants for analysis.

**Figure 1.**
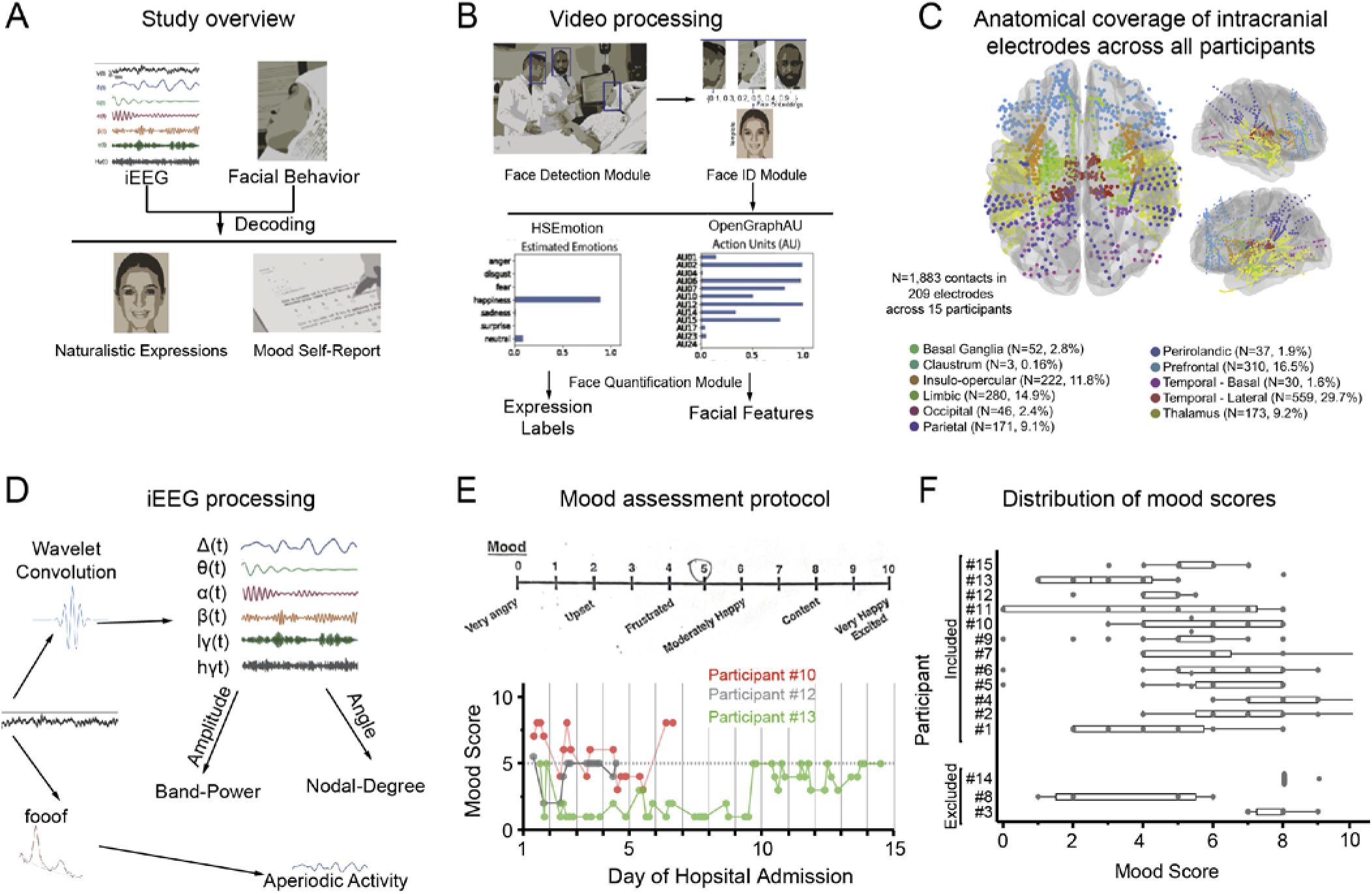
Multimodal neural and video recording paradigm of naturalistic facial expressions and self-reported mood. (A) Study overview showing simultaneous collection of continuous intracranial encephalography (iEEG) and video recordings of facial behavior from epilepsy patients during multi-day hospital stays. Periods of naturalistic smile and neutral facial expressions are labeled and participants periodically complete mood self-reports. iEEG and facial features are then used to decode expression labels and self-reported mood states. (B) Video processing pipeline using a custom, computer-vision pipeline which comprises three modules applied to inpatient video recordings. The face detection module identifies all faces per frame using MTCNN. The face ID module extracts facial embeddings using DeepFace and verifies identity against a participant template, enabling participant-specific tracking across frames. The face quantification module computes “emotion” probabilities using HSEmotion (2023) and facial action units (AUs) using OpenGraphAU (2022). These outputs are used in separate pipelines to generate candidate expression labels and facial features, respectively, used in decoders. (C) Anatomical coverage of intracranial electrodes across all participants, with electrode contacts color-coded by brain region of interest, demonstrating sampling across temporal, limbic, prefrontal, and other key areas involved in emotion and mood processing. (D) iEEG processing pipeline whereby recordings were preprocessed and analyzed in MATLAB. The broadband, aperiodic spectral slope (aperiodic activity, 1/f) between 30-80Hz was calculated using FOOOF, while band-limited power and nodal degree measures were extracted by time-frequency analysis using wavelet convolution. (E) Mood assessment protocol using an 11-point Likert scale (top) and mood scores recorded from 3 example participants over the course of their hospital stays (bottom). (F) Distribution of mood scores in all participants showing the spread in reported mood states across the hospital stay. Images representing individual faces in (A) and (B) are computer generated.

**Figure 2.**
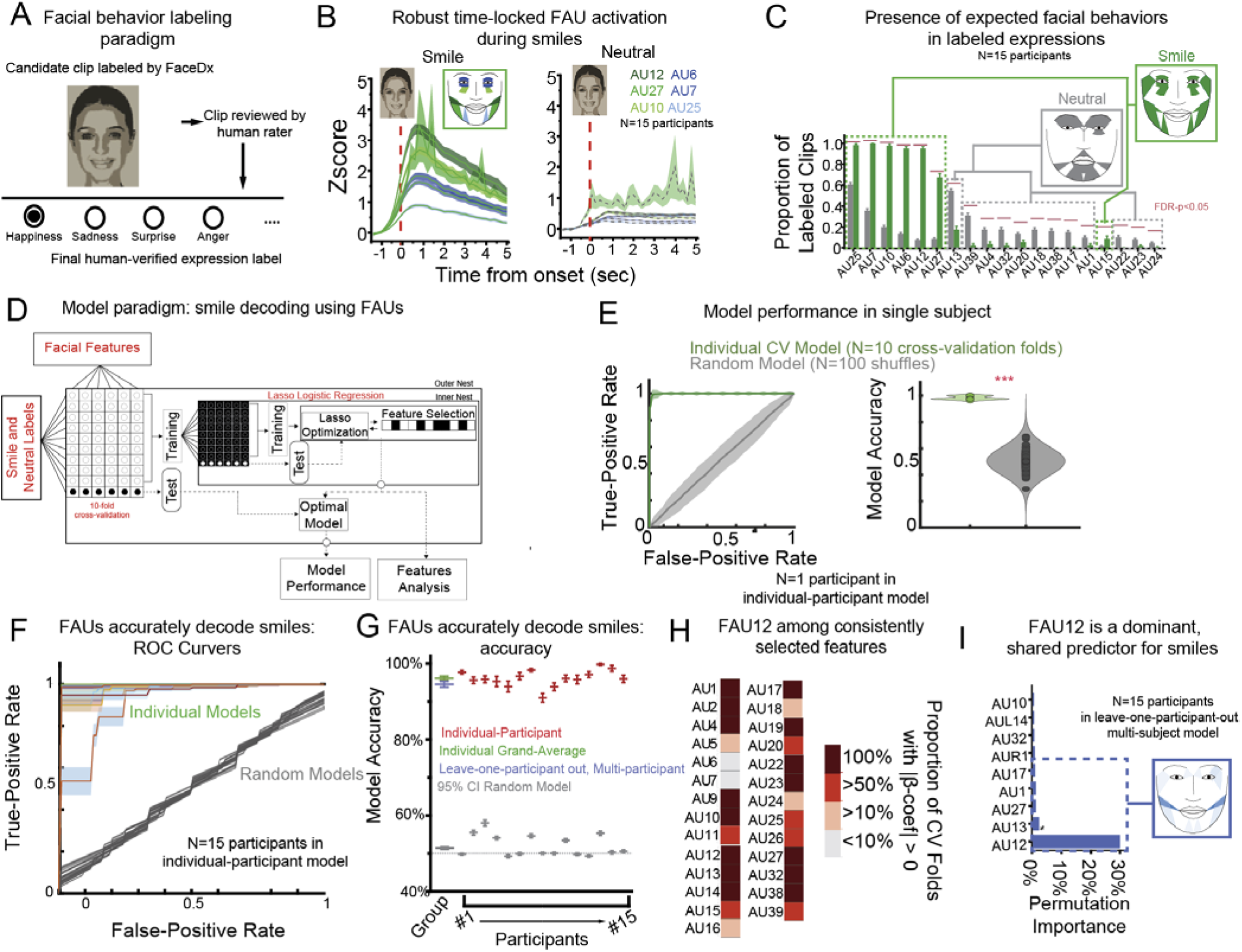
Automated video pipeline validates naturalistic smile detection with FAU12 as dominant behavioral signature. **(A) Facial Behavior Labeling Paradigm** in which independent raters reviewed candidate clips, epoched to expression onset and generated our automated behavioral pipeline, and labeled the expression **(B) Robust time-locked FAU activation during smiles** reveals increased activation of FAUs known to be associated with smile compared to neutral periods validates our detection pipeline. **(C) The presence of expected facial behaviors in labeled expressions**, in the bar plots, demonstrate key FAUs represented in facial recordings that distinguish smiles (green) from neutral expressions (gray). Simulated facial reconstructions illustrate the resulting expressions that align with expectations. **(D) Model Paradigm: Smile Decoding using FAUs** illustrates a model which takes in facial action units as features and expression labels as observations. Here, individual participant models employ <u>10-fold cross-validation</u> (CV), while multi-participant models use <u>leave-one-participant-out CV</u> to ensure generalizability. Inner loops using lasso <u>logistic regression</u> optimize regularization parameters and feature selection, while outer loops test performance on held-out data. **(E) Model performance in a single participant** validates the ability of the proposed generalizable model by demonstrating high performance across all individual-participant model CV folds (green) over a chance model (grey). Left: Mean receiver operating characteristics (ROC) curve of an example participant for model performance across cross-validation folds. Right: Accuracy across CV folds of the single model. **(F-G) FAUs accurately decode smiles** as demonstrated by high decoding performance for both individual-participant (F: colored, G: red) and multi-participant models (G: blue) compared to chance-level shuffled controls (Both: gray). **(H) FAU12 (lip corner elevation) is a dominant, shared (across participants) predictor for smiles** demonstrated by permutation importance of 30% (i.e., 30% drop in accuracy when the feature is randomly shuffled during modeling). Simulated facial expression demonstrates the key muscle activations underlying the top 5 features identified during multi-participant decoding. Images representing individual faces in (A) and (B) are computer generated.

To validate facial behavior labels, facial action units (FAUs) were extracted from labeled video in 4-second windows around each labeled event, starting 1 second prior to onset (Figure 2B). We observed robust activation in several FAUs known to be associated with smile over neutral expressions across all participants as illustrated in grand-average waveforms for FAU6 (i.e., cheek elevation), FAU25 (i.e., lip separation), FAU7 (i.e., lid tightener), FAU12 (i.e., lip corner elevation), FAU10 (i.e., upper lip elevation), and FAU27 (i.e., mouth stretch). Across participants, smiles were characterized by the presence of FAU6 (present in 97.5% of smile clips vs 40.9% of neutral clips), FAU25 (in 97.4% vs 66.2%), FAU7 (in 96.7 vs 43.8), FAU12 (in 93.6 vs 16.5), FAU10 (in 92.3 vs 22.4), and FAU27 (61.8 vs 14.3) (Figure 2C). FAUs with greater representation in neutral clips were heterogeneous and only present in a minority (<40%) of clips. Qualitatively, simulated facial expressions generated using the active FAUs during smile and neutral epochs aligned with expectations (Figure 2C insets).

Finally, to validate these labels in a naturalistic context, FAUs were used as features for lasso logistic regression models with a nested-, 10-fold cross-validation (CV) scheme^12,39^ to determine prediction accuracy on unseen testing data, independent from data used for training and model optimization (Figure 2D and 2E). This paradigm demonstrated high performance in decoding smiles using FAUs in individual participant models (Figure 2F, 2G; grand-average accuracy – green bar: 96.1%, standard error – i.e., S.E.:+/- 0.6%, p<0.01). Further, to assess generalization not only to new data from the same participant but also from an entirely new participant, a multi-participant model was constructed using a lasso logistic regression model with a nested-, leave-one-participant-out (LOPO) CV scheme whereby inner-fold training used data from all but one participant, and the outer-fold testing used unseen data from the left-out participant. FAUs could decode smiles with high accuracy in multi-participant, LOPO models (Figure 2G blue bar, mean accuracy: 94.5% S.E.:+/- 0.7%, p<0.01, 95% CI for accuracy of shuffled model: 50-51%). In multi-participant, LOPO CV modeling, up to 20 of the 41 FAUs were consistently selected (i.e., given a |ꞵ-coefficient| > 0 in the optimal model in >50% of CV folds; Figure 2H) but only 5 FAUs had permutation importance above 1% (Figure 2I; i.e., shuffling feature values associated with each FAU, individually and independently, in the optimal model and repeating prediction resulted in at least a 1% drop in accuracy). Again, qualitatively, a simulated facial expression using these FAUs aligned with expectations for a smile (Figure 2I right panel). FAU12 was the single most important feature for smile decoding (permutation importance: 30% vs the next highest of 2.4%). These findings corroborate knowledge of lip corner elevation (FAU12) as a dominant behavioral signature with high cross-participant generalizability.

### Facial expressions are robustly decoded from aperiodic activity in the temporal cortex across individuals

To determine if iEEG responses (Figure 1D) could decode facial expressions, iEEG data were epoched in 4-second windows around each labeled facial expression (Figure 2B), starting 1 second prior to onset, at each sampled node (i.e., Laplacian re-referenced channel on iEEG electrode; see Methods) for each spectral feature. iEEG features were fed into a nested, 10-fold CV lasso logistic regression model (Figure 3A, 3B). Spectro-spatial iEEG features could decode smiles with good accuracy in both individual participant models (Figure 3B, 3C red bar; grand-average accuracy - green bar: 85%, S.E.:+/-2.2%, p<0.01 for all individual participant models, grouped 95% CI accuracy of shuffled models: 48-61%) and in multi-participant, LOPO models (Figure 3D blue bar, mean accuracy: 79.5%, S.E.:+/-2.1%, p < 0.01, shuffled model 95% CI: 51 – 53%). To understand the neural dynamics underlying smile decoding, the iEEG features driving model performance were investigated. Aperiodic activity, high-gamma (hγ) power, and delta (δ) power in the lateral temporal cortex, along with lγ power in the thalamus were consistently selected in >50% of CV folds and in >50% of participants during individual modeling (Figure S2A) and selected in 100% CV folds in multi-participant, LOPO models (Figure 3E). In multi-participant modeling, the 3 most important (permutation importance >1%) were aperiodic activity in the lateral temporal cortex (permutation importance: 8.4%), hγ power in the temporal cortex (importance: 4.59%), and lγ power in the thalamus (importance: 1.91%) (Figure 3F; Figure S2B). The remaining features had permutation importance of <1%. To further assess feature importance, we repeated multi-participant modeling using only aperiodic activity in the temporal cortex and achieved statistically similar accuracy as when using all iEEG features (Figure 5B, green and red bars; mean accuracy: 74%, S.E.:+/- 3% vs 79.5%+/-2.1%, two-tailed t-test p > 0.05). In contrast, a model using a randomly selected feature with low importance (aperiodic activity in the thalamus) during smile decoding, performed statistically similar to random chance (Figure 5B, grey bar; mean accuracy: 53.4%, S.E.:+/- 3.8, two-tailed t-test p=0.12).

**Figure 3.**
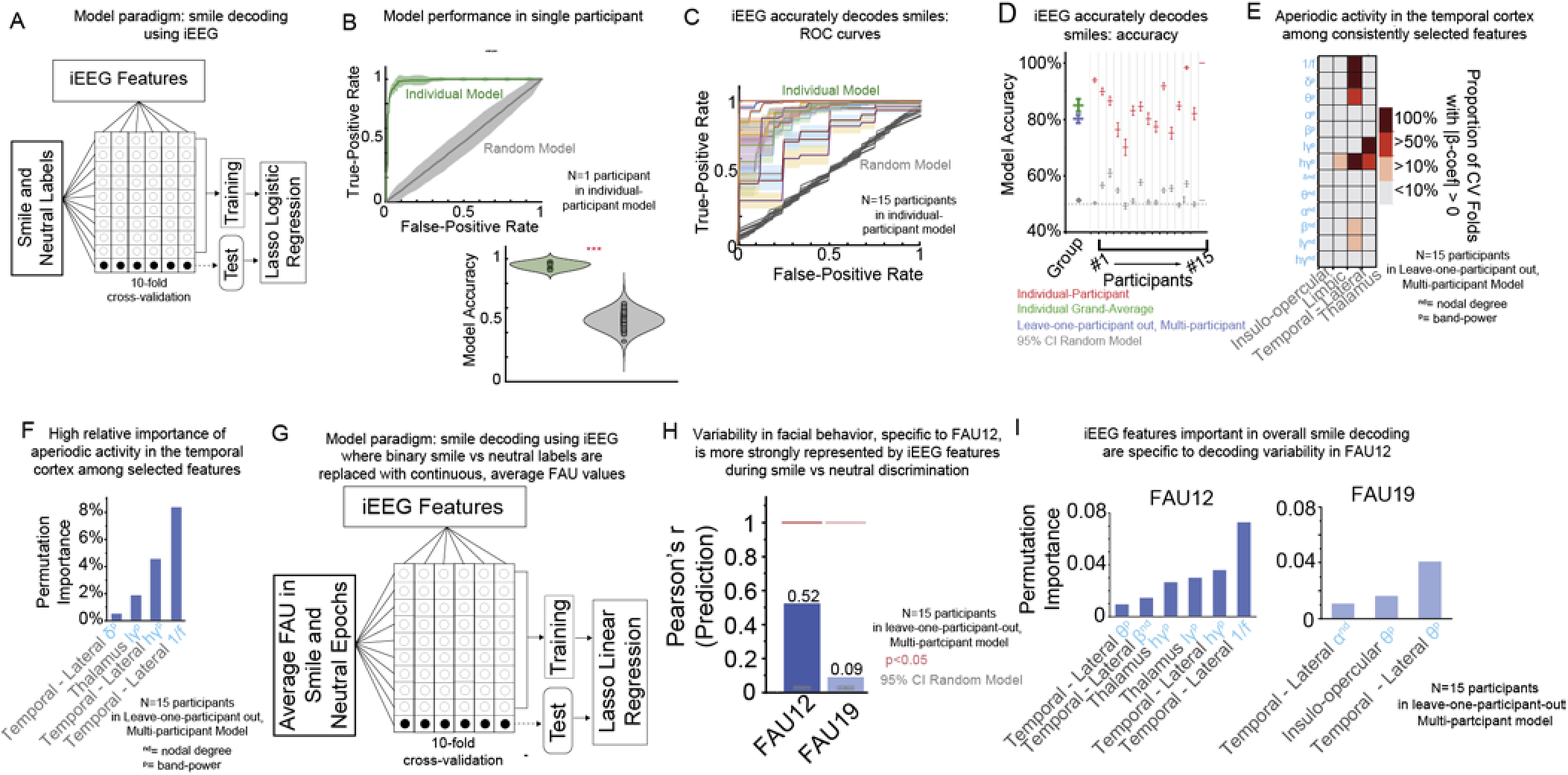
Aperiodic activity in the temporal cortex robustly decodes facial expressions across participants. **(A) Model Paradigm: Smile Decoding using iEEG** illustrates a model which takes in iEEG features and expression labels as observations. Here, individual participant models employ <u>10-fold cross-validation</u> (CV), while multi-participant models use <u>leave-one-participant-out CV</u> to ensure generalizability. Inner loops using lasso <u>logistic regression</u> optimize regularization parameters and feature selection, while outer loops test performance on held-out data. **(B) Model performance in a single participant** validates the ability of the proposed generalizable model by demonstrating high performance across CV folds (green) over a chance model (grey). Left: Mean receiver operating characteristics (ROC) curve of an example participant for model performance across cross-validation folds. Right: Accuracy across CV folds of the single model. **(C-D) iEEG accurately decodes smiles** as demonstrated by high decoding performance for both individual-participant (C: colored, D:red) and multi-participant models (D: blue) compared to chance-level shuffled controls (Both: gray). **(E) Aperiodic activity in the temporal cortex among consistently (>50% cross-validation folds) selected features** during multi-participant modeling. In addition, delta (δ) and high-gamma (hγ) power in the temporal cortex and low-gamma (lγ) power in the thalamus were consistently selected. **(F) High Relative Importance of aperiodic activity in the temporal cortex among selected features** as determined by permutation importance. This is followed by hγ power in the temporal cortex and lγ power in the thalamus**. (G) Model paradigm: smile decoding using iEEG where binary smile vs neutral labels are replaced with continuous, average FAU values.** Here, individual participant models employ <u>10-fold cross-validation</u> (CV), while multi-participant models use <u>leave-one-participant-out CV</u> to ensure generalizability. Inner loops using lasso <u>linear</u> <u>regression</u> optimize regularization parameters and feature selection, while outer loops test performance on held-out data. **(H) Variability in facial behavior, specific to FAU12, is more strongly represented by iEEG features during smile vs neutral discrimination**, to a larger magnitude over FAU19, which was significantly less important in decoding smiles. **(I) iEEG features important in overall smile decoding are specific to decoding variability in FAU12.** These features shared a similar distribution and magnitude while the set of features important in decoding FAU19 differ and have a lower overall magnitude of permutation importance.

Given the prominence of FAU12 characterizing and decoding smiles in our cohort, we next examined to what extent iEEG patterns of activation could directly and differentially encode specific facial patterns during facial expressions. iEEG features were modeled against FAU12 (i.e., an FAU with high importance during smile decoding, permutation importance: 30%) and FAU19 (i.e., an FAU with low importance during smile decoding, permutation importance: 0.3%) (Figure 3G) by taking the FAU value at the onset of the expression event. Modeling was completed using a nested, 10-fold CV, lasso linear regression model. A multi-participant, LOPO model using all iEEG spectro-spatial features to decode FAU12 achieved good performance (Figure 3H; Pearson’s r:0.52, p<0.01, shuffled model Pearson’s r 95% CI: 0.01-0.03) compared to lower performance with decoding FAU19 (Pearson’s r: 0.09, p<0.01, shuffled model Pearson’s r 95% CI : 0.02-0.04). To decode FAU12, aperiodic activity in the lateral temporal cortex was the most important feature (permutation importance: 0.07, i.e., a decrease of 0.07 in the Pearson’s r when the feature was shuffled). Further, the top 3 features in linear decoding of FAU12 were identical to the top 3 identified in overall binary expression label decoding (Figure 3I, left panel). In contrast, decoding FAU19 relied on a different, non-overlapping distribution of iEEG features with lower overall permutation importance (Figure 3I, right panel).

In summary, aperiodic activity in the lateral temporal cortex emerged as the dominant neural signature for smile decoding, demonstrating precise neural-behavioral correspondence by specifically encoding the key facial action unit (FAU12) that defines smiling behavior.

### Mood decoding succeeds only in individuals with limbic γ power encoding mood states

To assay longitudinal trends in mood, participants completed a brief survey question (on an 11-point Likert scale: 0-very angry to 10–very happy) periodically during their hospital stay (Figure 1A, 1E). Of the 15 participants meeting criteria for smile decoding (see Methods), 12 completed sufficient mood surveys for inclusion (12-70 surveys per participant, average range of scores: 6.12, average number of unique scores: 5.67, Figure 1F, Table S1). Of the 15 participants, 3 were excluded -- 1 for repeatedly reporting the same 1 or 2 scores, and 2 for completing only 5 surveys. To determine if mood scores correlated with validated mood measures, we compared post-operative mood scores to Beck Depression Inventory^40^ (BDI) scores obtained pre-operatively as part of a standard clinical assessment battery. At the group level, there were no statistically significant linear relationships between pre-operative BDI and post-operative individual mood scores (Pearson’s r:-0.043, two-tailed t-test p=0.47), mood scores averaged across each patient’s stay (Pearson’s r:0.079, two-tailed t-test p=0.8), or with the first reported mood scores (which are more temporally proximal to pre-operative BDI scoring, Pearson’s r:-0.35, two-tailed t-test p=0.25).

We next tested if FAUs and iEEG could decode mood scores. To ensure stability and consistency over extended periods of time, FAUs and iEEG were averaged in 6 incremental time blocks (5, 10, 15, 20, 25, and 30 minutes) centered at onset of survey completion (Figure 4A). Features were fed into a lasso linear regression model with a nested-, leave-one-out (LOO) CV scheme for individual-participant models (Figure 4B and 4C). A feature was deemed consistent if it demonstrated significant decoding performance in greater than 3 of the 6 time blocks (>50%). FAUs failed to significantly and consistently decode mood (in >50% of time blocks) in all but 2 of the 12 participants (Figure S3A). In these 2 participants, there was no shared, consistent distribution of FAUs underlying decoding. To further probe the relationship between facial features and mood, we expanded our analysis to include a broader range of time windows (30 to 240 minutes) and an extended feature set incorporating multiple facial analysis toolkits, including OpenFace^41^. Across these conditions, facial expression features alone failed to consistently predict mood in the majority of participants, with no statistically significant decoding at the group level (Figure S5).

**Figure 4.**
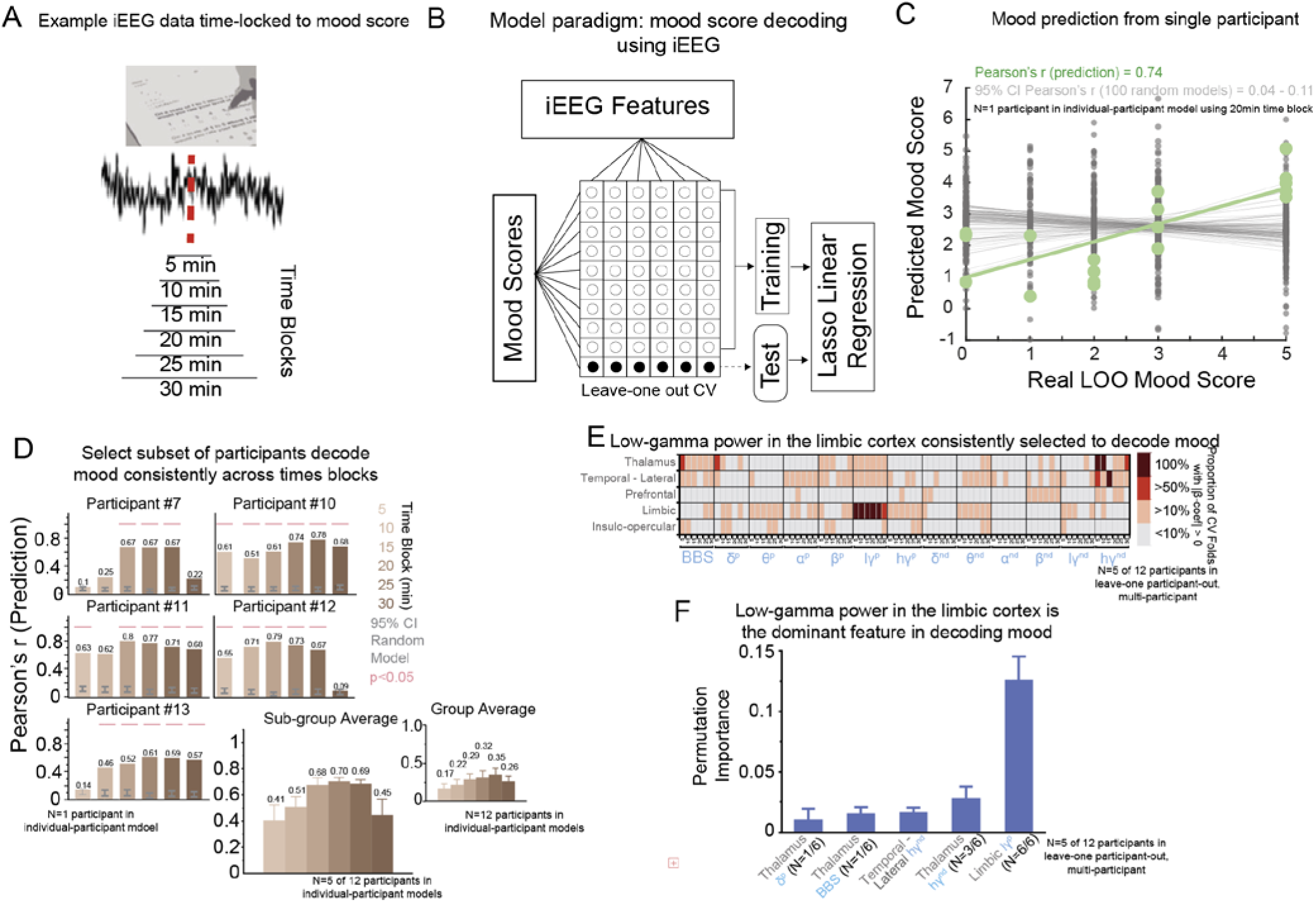
Limbic gamma activity predicts mood in subset of individuals. **(A) Example iEEG Data Time-locked to Mood Score** depicting the 6 time blocks used in analysis ranging from 5 to 30 minutes around onset of mood survey completion. **(B) Model Paradigm: Mood Score Decoding using iEEG** illustrates a model which takes in iEEG features and mood scores as observations. Here, individual participant models employ <u>leave-one-out cross-validation</u> (CV), while multi-participant models use <u>leave-one-participant-out CV</u> to ensure generalizability. Inner loops using lasso <u>linear regression</u> optimize regularization parameters and feature selection, while outer loops test performance on held-out data**. (C) Mood Prediction from a Single Patient** validates the ability of the proposed generalizable model by demonstrating high performance across in a single individual-participant model CV folds (green) over a chance model (grey). **(D) Select Subset of Participants Decode Mood Consistently Across Time Blocks** (>50% of time blocks) as demonstrated by high predictive Pearson’s r (bar plots), compared to random models (grey error bar). Sub-group average (inset) of participants with consistent decoding. **(E) Low-gamma (lγ) Power in the Limbic Cortex Consistently Selected to Decode Mood** (>50% cross-validation folds) across all 6 time blocks during multi-participant modeling. **(F) lγ power in the limbic cortex is the dominant feature in decoding mood** compared to all other features as determined by permutation importance. This is followed by high-gamma (hγ) nodal degree in the thalamus and lateral temporal cortex.

**Figure 5.**
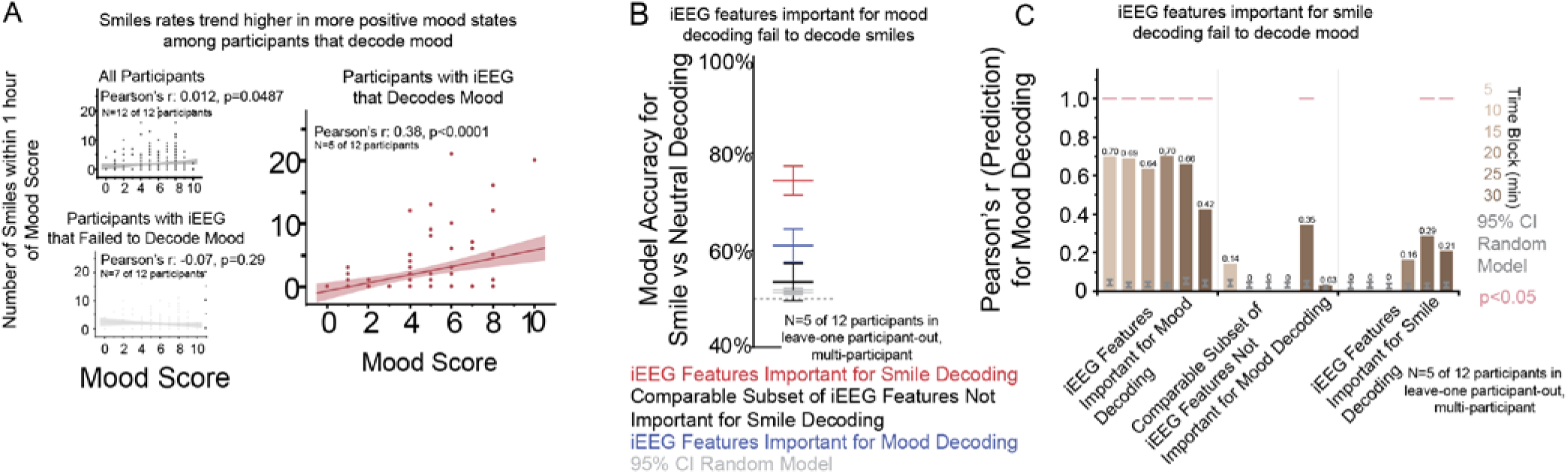
Neural features encoding naturalistic facial expressions are dissociated from mood decoding. **(A) Smiles Rates Trend Higher in More Positive Mood States Among Participants that Decode Mood** (right panel) and this trend was not observed among participants without consistently decodable mood using iEEG (left panel). **(B) Mood-related Features Fail to Decode Smiles** (blue bar), among the 5 participants with consistent mood decoding, performing similarly to iEEG features that had low importance for smile decoding.

Of the 12 participants with sufficient mood scores, 5 participants showed significant, consistent mood decoding using iEEG (Figure 4C, 4D, Pearson’s r range of means across time blocks: 0.41-0.70, shuffled model 95% CI: 0.09 – 0.15; Figure S3B). Compared to the remaining participants without decoding, these 5 participants showed no differences in the range of mood scores reported (two-tailed t(12.2)=-0.16, p=0.87), total number of mood scores completed (two-tailed t(9.5)=0.27, p=0.79), or number of unique mood scores reported (two-tailed t(11.5)=-0.38, p=0.71). In individual decoding of these 5 participants, there was no consistently shared distribution of iEEG spectro-spatial features contributing to performance across multiple time blocks. In multi-participant, LOPO modeling using these 5 participants, lγ power in limbic regions was the most salient feature selected in >50% CV folds in all time windows (Figure 4El), and the feature with the highest importance (Figure 4F; average permutation importance: 0.13, S.E.:+/-0.02). When used as the sole feature in multi-participant modeling, lγ power in limbic regions performed well across all time blocks (Figure 5C left panel, Pearson’s r range: 0.42 – 0.70, p<0.01). Repeating this analysis with a randomly selected feature with low permutation importance during mood decoding (lγ power in insulo-opercular regions) resulted in poor performance with iEEG in 5 of the 6 time blocks performing statistically similar to random chance (Figure 5C middle panel, p > 0.05). In summary, mood decoding from neural signals succeeded only in a subset of participants, with low-gamma (lγ) power in limbic regions serving as an important neural signature in those with decodable mood states.

### Expression and mood neural features are dissociated

Despite the intuitive relationship between facial expressions and mood, cross-domain analysis revealed neural dissociation between their encoding mechanisms. To investigate if the frequency of naturalistic smiles correlated with mood scores, the number of smile episodes occurring within a 2-hour window centered at completion of each survey were tallied. At the group level, there was no statistically significant correlation between the number of smiles and individual mood scores (Figure 5A top left; Pearson’s r:-0.07, two-tailed t-test p=0.29). To assess if there was a relationship with smile counts and participant specific mood ranges, mood scores were split into high and low scores based on individual participant median split. There was no statistically significant correlation between the number of smiles and the binarized mood state (two-tailed t(210)=-0.86, p=0.39). Finally, there was no correlation between mood scores, and smile counts when averaged over the entire hospital at the group level (Pearson’s r:-0.07, two-tailed t-test p=0.80).

Given the heterogeneity in individual participant models using iEEG to decode mood, we repeated analysis on smile counts stratified by ability to decode mood (Figure 3A right). For the 5 participants with models that were able to decode mood using iEEG, happier mood was associated with increased smile counts in the 2 hours surrounding survey completion (Pearson’s r: 0.38, p < 0.0001). No significant correlation was seen in the remaining participants in whom iEEG failed to decode mood (Figure 5A right bottom; Pearson’s r: -0.07, p=0.29). Correlations between mood and BDI remained non-significant after a similar stratification (two-tailed t-test p>0.1).

Finally, we assessed whether iEEG features important to smile decoding could decode mood and if those for mood could decode smiles. lγ power in limbic regions, a feature with high permutation importance in mood decoding, was used as the only feature during smile decoding and performed poorly (Figure 5B; average accuracy: 61%, S.E.:+/-0.04). This performance was not statistically different from a randomly selected feature which had low permutation importance in smile decoding (two-tailed t-test p>0.05). Aperiodic activity in the temporal cortex, a feature with high permutation importance in smile decoding, was used as the sole feature during mood decoding and performed poorly (Figure 5C right panel, performance in 4 of the 6 time blocks was statistically similar to chance, p > 0.05).

In summary, expression-mood behavioral relationships emerged in participants with decodable neural mood signatures, and cross-domain analysis revealed neural dissociation with features optimized for expression decoding failing to predict mood and vice versa, suggesting distinct neural mechanisms.

## Discussion

This study provides the first investigation using simultaneous intracranial electroencephalography (iEEG) and facial expression monitoring to decode both naturalistic smiles and self-reported mood states from the same neural signals over multiple days. Given the intuitive behavioral relationship between facial expressions and mood, we hypothesized that their neural encoding mechanisms would share common substrates and demonstrate overlapping patterns of brain activity. Using continuous iEEG recordings from 15 participants, we captured 1,396 smiles and 3,746 neutral expressions alongside periodic mood assessments over multi-day hospital stays. We demonstrated five key findings: 1) Naturalistic smiles were decoded with high accuracy using both facial action units (94.5% accuracy) and iEEG signals (79.5% accuracy), with aperiodic activity in the lateral temporal cortex serving as the most predictive neural feature. 2) The key facial feature for smile detection (lip corner elevation, FAU12) was specifically and reliably encoded by the same temporal cortex neural signals across individuals. 3) In contrast to the robust and consistent smile decoding results, mood prediction from neural signals succeeded only in a subset of participants (5 of 12), with lγ power in limbic regions as the key distinguishing feature. 4) Expression-mood behavioral relationships emerged exclusively in participants with decodable neural mood signatures – those with successful neural mood decoding showed correlations between smile frequency and self-reported mood (r=0.38), while others showed no relationship. 5) Neural dissociation was observed between expression and mood encoding: features optimized for smile decoding failed to predict mood, and vice versa, indicating distinct non-overlapping mechanisms. Together, these findings challenge assumptions about expression-mood neural relationships, establish robust mechanisms for facial expression processing while revealing complex individual differences in mood encoding, and provide a foundation for developing personalized biomarkers in precision psychiatry.

### Temporal cortex as a robust neural substrate for facial expression processing

Our findings establish aperiodic activity in the lateral temporal cortex as a remarkably consistent and generalizable neural signature underlying facial expression decoding across individuals. The ventral and lateral temporal regions have long been recognized as critical nodes for biological motion perception and social cognition, with neuroimaging studies consistently demonstrating activation in this area during facial expression processing^42–44^. Increased activation of the middle temporal gyrus, among various motor and limbic regions, has been shown to start seconds prior to laughter onset and linked to laughter detection^45^. Additionally, activation of superior and medial temporal regions are shared across individuals during both observation of and execution of happy facial expressions over non-emotional facial expression^46^. Further, electrical cortical recordings of temporal regions have been shown to be involved in mirth and laughter in various case reports, with some inducing these behaviors through direct stimulation^47–49^. Finally, studies have linked perturbations in activity among regions involving the middle temporal gyrus in syndromes such as pathological laughing and crying, which are characterized by abnormal emotional expressions^50^. Building on this foundation, our multi-participant models demonstrated that aperiodic activity in the lateral temporal cortex was selected in 100% of cross-validation folds and carried the highest permutation importance (8.4%) for smile decoding (Figure 2D). Remarkably, this same neural feature specifically encoded the most behaviorally relevant facial action unit-lip corner elevation (FAU12) – which itself carried 30% permutation importance in facial decoding models (Figure 2E). When aperiodic activity in the temporal cortex was used as the sole neural feature, it achieved statistically comparable performance (74% accuracy) to models using the full feature set (79.5% accuracy), underscoring its singular importance. This precise neural-behavioral correspondence, where the same temporal cortex signals that decode overall smiling behavior is also associated with the key muscular component of smiling, suggests a remarkably organized and consistent neural representation. These findings align with and extend the existing literature by demonstrating that temporal cortex aperiodic activity serves as a reliable neural readout for naturalistic facial expressions recorded over multiple days in real-world settings. Unlike previous studies that relied on controlled laboratory presentations of static facial images or brief video clips, our approach captured spontaneous expressions across diverse naturalistic contexts – with visitors, watching television, using personal devices – yet still yielded highly consistent neural signatures. In summary, aperiodic activity in lateral temporal cortex represents a robust, generalizable neural substrate for facial expression processing that maintains precision at both the behavioral level (overall smile detection) and the granular muscular level (specific facial action units), establishing this region as potential neural readout for naturalistic social emotional communication.

### Limbic γ power and the neural heterogeneity of mood encoding

The striking individual heterogeneity observed in mood decoding reveals differences in the neural encoding of mood states across individuals, with lγ power in limbic regions emerging as a critical but variable neural signature. Gamma power in limbic structures has been implicated in mood regulation in human intracranial studies, demonstrating that γ rhythms serve as biomarkers for major depression^51^, distinguish individuals with mood disorders from healthy controls^40^,f and decode depression severity^52^. Building on this foundation, our finding that only 5 of 12 participants showed consistent mood decoding through lγ power in limbic regions (Figure 3B) suggests that mood encoding mechanisms are not universally implemented across individuals. Interestingly, this effect only appears during multi-subject modeling, and not individual subjects modeling, which may highlight the limits of the low number and variability of mood score observations in individual participants. The participants with successful neural mood prediction demonstrated robust correlations between limbic γ activity and self-reported mood states (r=0.42-0.70), while those without decodable mood signatures showed no such relationships, indicating differences in limbic network dynamics. This heterogeneity may reflect individual differences in underlying limbic circuit psychopathology, including different epileptogenic networks, or inherent variability in the neural representation of sustained affective states. Recent large-scale human intracranial studies have similarly revealed that mood-predictive networks vary substantially across individuals, with some participants showing amygdala-hippocampus subnetworks that encode mood variations while others lack these coordinated patterns entirely^53^. The observation that expression-mood behavioral correlations emerged exclusively in participants with decodable neural mood signatures further supports the hypothesis that limbic γ power may serve as a potential biomarker linking momentary expressions with sustained mood states. In summary, lγ power in limbic regions represents an individually variable neural mechanism for mood encoding, suggesting that mood disorders may involve heterogeneous disruptions of limbic power dynamics that require personalized approaches to neural biomarker development and therapeutic intervention.

### Neural dissociation reveals distinct mechanisms for momentary expression and sustained affective states

The neural dissociation between expression and mood encoding mechanisms reveals distinct temporal architectures optimized for different aspects of affective processing, challenging assumptions about shared neural substrates for facial expressions and internal emotional states^22–27^. Facial expression processing operates on rapid timescales, with neural responses to emotional faces beginning as early as 130 milliseconds post-stimulus and reflecting fast, stimulus-driven computations designed for immediate social communication^42^. s stimulus^23,24,54^. These rapid responses align with the distributed cortical networks identified across species, where temporal cortex neurons respond selectively to dynamic facial features within hundreds of milliseconds, enabling real-time social signal detection^43^. Our cross-domain analysis demonstrated that aperiodic activity in the temporal cortex – optimized for these rapid, discrete expression events – failed to predict mood states, while limbic γ power critical for mood encoding showed no relationship to smile detection. This dissociation may reflect the difference between transient, stimulus-locked neural responses that support moment-to-moment social signaling versus sustained network configurations that encode persistent internal states. Mood states, unlike discrete expressions, emerge from distributed limbic-prefrontal networks operating over extended timescales of minutes to hours, requiring sustained patterns of neural activity rather than rapid stimulus-response dynamics^3,19^. The temporal mismatch between these systems provides a mechanistic explanation for why neural features excelling at one domain fail in the other. The observed neural dissociation may also reflect evolutionary optimization, where rapid expression detection serves immediate survival needs through fast social communication, while sustained mood networks integrate complex environmental and internal signals over extended periods to guide longer-term behavioral strategies. Further, while smiles, operationalized by activation of FAU12 (lip corner puller), are a canonical marker of positive affect in behavioral science, individuals can display smiles despite dysphoria or suppress overt affect despite euthymia^2^. In summary, the lack of cross-domain neural transferability reveals that momentary facial expressions and sustained mood states are likely supported by distinct neural mechanisms operating on distinct temporal scales, suggesting that care should be taken when developing objective biomarkers for mood disorders to target mood-specific neural signatures rather than expression-derived proxies.

### Methodological innovations and clinical translation

This study introduces several methodological advances that enhance the feasibility and precision of naturalistic neurobehavioral research. The video-facial behavior pipeline represents a significant improvement in automated, markerless facial expression analysis for clinical settings, combining real-time face detection, identity verification, and multi-modal feature extraction in a unified framework optimized for extended, uncontrolled recordings. Unlike laboratory-based paradigms that rely on posed expressions or brief recording sessions, our pipeline enables continuous behavioral quantification across multiple days of naturalistic behavior, dramatically expanding the ecological validity and statistical power of neurobehavioral studies. The semi-automated labeling approach, which uses computer vision for initial candidate identification followed by manual verification, achieved efficient processing of over a week of continuous recordings per participant, a scale impractical with manual annotation methods. This hybrid approach maintains high labeling accuracy while reducing the labor-intensive bottleneck that has historically limited naturalistic studies to small datasets or brief recording periods. Additionally, our implementation of simultaneous high-resolution iEEG and facial expression monitoring over extended periods provides unprecedented temporal precision for investigating neural-behavioral relationships, bridging the gap between controlled laboratory paradigms and real-world clinical applications. These methodological innovations collectively enable scalable, objective biomarker development for psychiatric conditions while maintaining the nuanced behavioral context essential for understanding individual differences in neural encoding. The clinical translation potential is particularly notable, as these approaches could be adapted for routine monitoring in care settings, providing objective measures to complement traditional subjective assessments and potentially enabling personalized treatment optimization based on individual neural-behavioral signatures.

### Limitations

This study has several important limitations that constrain interpretation of the findings. First, our study utilized epilepsy patients with implanted electrodes, whose brain networks may differ from healthy populations due to underlying pathophysiology and medication effects. The clinical setting and relatively sparse limbic electrode coverage, driven by clinical rather than research considerations, may have limited our ability to fully characterize mood-related networks. Second, our mood assessment relied on a simplified 11-point Likert scale that was chosen for feasibility in obtaining repeated measurements during the clinical stay, though this approach conflates valence and arousal dimensions. This scale did not correlate with established depression scales (BDI), raising questions about construct validity. However, it is worth noting that these measures assess different constructs and timeframes – the BDI captures depressive symptoms over the preceding week and was administered preoperatively, while our Likert scale assessed momentary mood states during the hospital stay – making lack of direct correlation potentially not indicative of construct validity issues. Regardless, more comprehensive mood assessment tools with validated psychometric properties would strengthen future investigations. Third, the temporal mismatch between brief facial expressions (4-second epochs) and extended mood assessments (5-30 minute windows) may obscure meaningful relationships between these phenomena. Fourth, our relatively small sample size (N=15 for expression analysis, N=12 for mood analysis) limits statistical power and generalizability, particularly given the individual heterogeneity observed in mood decoding. Fifth, we employed linear regression models rather than more sophisticated machine learning approaches that might better capture complex nonlinear relationships between neural features and behavioral outcomes. Finally, the in-room video quality and naturalistic recording conditions, while ecologically valid, may have introduced variability that more controlled laboratory settings could minimize.

## Methods & Materials

Multimodal, intracranial electroencephalography (iEEG) and video-facial recordings over several days were obtained from 16 epilepsy participants (Figure 1A, Table S1; 7 females, age range: 24-59 years old) undergoing inpatient seizure monitoring. As detailed below, 15 participants met inclusion criteria for behavioral expression decoding and 12 participants met inclusion criteria for mood decoding. All participants provided informed consent, monitored by the Stanford University Institutional Review Board, in accordance with the ethical standards of the Declaration of Helsinki. The decisions regarding electrode implantation, targets, and duration were made entirely based on clinical grounds, independent of this investigation. Participants were informed that their involvement in the study would not affect their clinical treatment, and they could withdraw at any time without compromising their care.

### Smile Behavioral Labeling and Data Segmentation

All participants in the epilepsy monitoring unit were routinely recorded with high-definition audio at 48 kHz and RGB video (640 x 480 pixels at 30 frames per second) 24 hours a day. The camera, located on the room ceiling, was manually centered on the participants’ faces by video technologists throughout their stay. Our methods for extracting facial dynamics are previously described^12^. Briefly, video was down-sampled to 5 frames per second and in each frame, the participant’s face was identified and isolated from all faces present (MTCNN^55^ and DeepFace^56^; Figure 1B). Hundreds of candidate smile- and neutral-facial expressions in the multi-day video recordings were filtered using a facial emotion model (HSEmotion 2023^57^) to make inferences and pre-label candidate ‘happiness’ and ‘neutral’ periods which were then extracted in 4 second epochs starting 1 second prior to event onset. Candidate events were manually reviewed by two independent human raters blinded to the automated labels and to each other’s labeling. Only events with consensus labeling between raters were selected for processing. Manual raters were instructed to only label faces with unobstructed views. Events occurring during periods involving behavior that would obscure an expression (e.g., eating or brushing teeth) were also excluded. Neutral facial expressions were defined as having no perceived emotional expression (i.e., ‘blank face’). Multi-day iEEG and visual-facial recordings used in subsequent analyses were segmented around these final labels and used as features in our facial expression decoder.

### Facial dynamics data acquisition

To investigate behavioral changes during any given period of time, frame-by-frame facial action units (FAUs) were extracted using a custom computer vision pipeline (Figure 1C). Videos were downsampled to 5 frames per second, with faces identified in each frame via MTCNN. Frames without faces were skipped, and those with multiple faces underwent facial verification via DeepFace using a pre-trained VGG-Face model to extract facial embeddings and match them with the provided participant image. As part of this pipeline, a custom-built algorithm tracked face displacement between frames, only triggering verification when distance exceeded a predefined threshold. Videos were automatically cropped to include only the verified participant’s face and processed through OpenGraphAU^58^, an open-source transformer for AU detection, as well as HSEmotion - an open-source convolutional neural network (CNN) for “emotion” recognition. This generated frame-by-frame outputs for 41 AUs (AU1, AU2, AU4, AU5, AU6, AU7, AU9, AU10, AU11, AU12, AU13, AU14, AU15, AU16, AU17, AU18, AU19, AU20, AU22, AU23, AU24, AU25, AU26, AU27, AU32, AU38, AU39, AUL1, AUR1, AUL2, UR2, AUL4, AUR4, AUL6, AUR6, AUL10, AUR10, AUL12, AUR12, AUL14, and AUR14) as well as 7 emotions (Happiness, Sadness, Surprise, Fear, Anger, Disgust, and Contempt).

For behavioral analysis, raw FAU intensity values were converted into binary indicators. Further continuous emotion probabilities were calculated, binarized, and segmented to identify sequences of consecutive frames with detected emotions. Short-term decoding involved manual labeling of approximately 100-300 instances each of smile and neutral, with our automated pipeline filtering the search space to approximately 100-500 clips per emotion per patient for efficient manual review (Figure 2A).

### Mood Surveys

We employed a custom, simple, one question survey in which participants were asked to rate their mood on an 11-point Likert scale (from 0: Very Angry to 10: Happy/Content) at 2-3 intervals each day for the duration of their clinical EMU stay (Figure 1E; Figure S1). Inclusion criteria were created to optimize variance and sample size by requiring at least 3 unique scores and a distribution of scores providing greater than 10^3^ unique permutations. Participants were provided with an iPad which contained a link to the survey and that provided reminders throughout the day. Multi-day iEEG and visual-facial recordings used in subsequent analyses were segmented around the completion time of each survey and used as features in our decoder. Given that mood metrics are expected to be stable over extended periods of time, mood-associated analysis was performed independently on data in 6 different time blocks; 5, 10, 15, 20, 25, and 30 minutes before and after survey completion time.

### Electrode registration and anatomical parcellation

Electrode location (Adtech Medical; centre-to-centre contact spacing of 3 mm; N=2,037 contacts across 219 electrodes) in 3D space was obtained from post-implant CT co-registered with the participant’s pre-operative MRI. The anatomic location of each contact was determined using FreeSurfer-based^59^ automated parcellation, as described previously^12^, based on the ‘FSLabel_aparc_aseg’ atlas^60^. Electrode contacts were further anatomically categorized into the following regions of interest (ROI, Figure 1C): basal ganglia (N=53 contacts in 13/15 participants), claustrum (N=3 contacts in 2/15 participants), insulo-opercular cortex (N=253 contacts in 15/15 participants), limbic (N=303 contacts in 15/15 participants), occipital (N=49 contacts in 6/15 participants), parietal (N=213 contacts in 14/15 participants), perirolandic (N=42 contacts in 6/15 participants), prefrontal (N=336 contacts in 13/15 participants), basal temporal cortex (N=30 contacts in 8/15 participants), lateral temporal cortex (N=571 contacts in 15/15 participants), and thalamus (N=184 contacts in 15/15 participants). Limbic nodes included contacts in the amygdala, cingulate, and hippocampus. Contacts located outside of the brain or in lesional tissue were excluded. For electrode visualization, the FreeSurfer average brain was used with coordinates in standard Montreal Neurologic Institute (MNI) space.

### iEEG Data acquisition and signal preprocessing

iEEG recordings were sampled at 1024Hz (Nihon Koden) and preprocessed and analyzed in MATLAB using the FieldTrip toolbox^61^, FOOOF toolbox^62^, and custom scripts. Preprocessing consisted of a fourth-order notch filter to attenuate line noise (60, 120, and 180Hz), and Laplacian re-referencing to minimize far-field volume conduction. The broadband, aperiodic spectral slope (aperiodic activity, 1/f) between 30-80Hz was calculated using FOOOF. Separately, the signal was passed through 8^th^-order zero-phase IIR Butterworth filters followed by Hilbert transform within standard bands: delta (δ; 1–4Hz), theta (θ; 4–8Hz), alpha (α; 8–12Hz), beta (β; 15–25Hz), low-gamma (lγ; 25–70Hz) and high-gamma (hγ; 70–170Hz). Band power was extracted as the squared absolute value of the resultant complex signal. Band phase was extracted as the angle of the Hilbert transform from which we calculated inter-site phase clustering between all node pairs with a threshold of 1 standard deviation above the median^63^. Nodal degree was calculated as the number of suprathreshold nodal pairs associated with a respective node in each band.

### iEEG and Facial Feature Modeling to Predict Smiles

To investigate if iEEG activity could predict between smiles and neutral events, logistic regression with L1 regularization and repeated (3 times) nested 10-fold cross-validation (CV) was employed (Figure 2A). Observation classes were balanced by selecting all smile events and randomly selecting an equal number of neutral events in each participant. Individual participant regression models were independently trained with either iEEG spectro-spatial features or FAUs. With models using iEEG features, each feature comprised a spectro-spatial pair of a node (i.e., Laplacian re-referenced channel on an electrode) and a given spectral signal (e.g., average θ power). The spectral signal was averaged over a 4 second epoch for each observation. As part of the CV scheme, in the inner fold, the regularization strength was optimized and in turn, the features set was filtered, using 10-fold CV on the training data made up of 9 of the 10 outer-CV folds. To avoid data leakage and optimize model performance, features were normalized after the CV fold split and only within the inner fold. The optimized model was then tested on the left-out data from the outer-CV fold to generate accuracy measures. This process was replicated 10 times for each outer fold and repeated 3 times with different outer-nest, CV-fold splits to ensure stability. This generated a distribution of 30 independent measures of accuracy and sets of selected features.

Given the clinical importance of generalizable models that perform well across different participants, multi-participant, leave-one-participant-out (LOPO) models were generated using the previously described regression model scheme. Given the limitations of logistic regression in the setting of features with a heterogeneous number and positioning of sampled iEEG nodes, multi-participant modeling was limited to ROIs shared by all participants (insulo-opercular cortex, limbic, lateral temporal cortex, and thalamus). All participants had at least 3 nodes in each shared ROI. In each participant, all the nodes within an ROI for a given spectral signal were grouped and partial least squares regression was implemented to order nodes within an ROI by degree of covariance with the predictor. The top 2 nodes in each ROI were selected and grouped together across participants. For the CV scheme, nested leave-one-participant-out CV was performed whereby N-1 participant datasets were used for training in the inner fold and tested against the left-out participant dataset in the outer fold.

### iEEG and Facial Feature Modeling to Predict Mood

To investigate if iEEG activity could predict mood, linear regression with L1 regularization and nested LOO CV was employed (Figure 2A). Independent models were created for each of the 6 time blocks to test stability. The spectral signal was averaged in each of the 6 time blocks for each observation. Inner fold optimization was performed using LOO CV on the training data from the k-1 observations in the outer fold. To avoid data leakage and optimize model performance, features were normalized after the CV fold split and only within the inner fold. The optimized model was then tested on the left-out observation from the outer-CV fold. The distribution of predictions from the k LOO fold iterations was then compared to the actual left-out observations by using Pearson’s r to quantify correlation and performance. Multi-participant modeling was performed as previously described.

### Statistical Analysis

In both single- and multi-participant models, the CV model performance (average accuracy for smile decoding, and Pearson’s r for mood decoding), and the proportion of times each feature was selected across all CV folds were calculated. Statistical significance (α=0.05) of the model’s performance was performed by repeating all modeling steps with random permutations of observations (n=100; shuffled model). Feature salience was reported at the ROI-spectral signal level. An ROI-level feature (e.g., δ power in limbic regions) was considered salient if at least one spectro-spatial signal feature (e.g., θ power in node x) within the ROI was selected in greater than 50% of CV folds, in greater than 50% of participants. Further, salient ROI-level features were ordered by permutation importance whereby each individual feature in the testing data was randomly permuted independently and prediction repeated to determine if and how much performance changed. This was repeated 100 times with different random permutations to ensure stability.

## Acknowledgements

We extend gratitude to all our research participants. We would also like to acknowledge the generous contributions of the members of the Precision Neurotherapeutics Laboratory for helpful feedback on the manuscript and throughout the course of the study.

## Data and Code Availability Statement

The data that support the findings of this study are available from the corresponding author upon reasonable request, subject to appropriate ethics approval and data use agreements.

---

**Figure S1.**
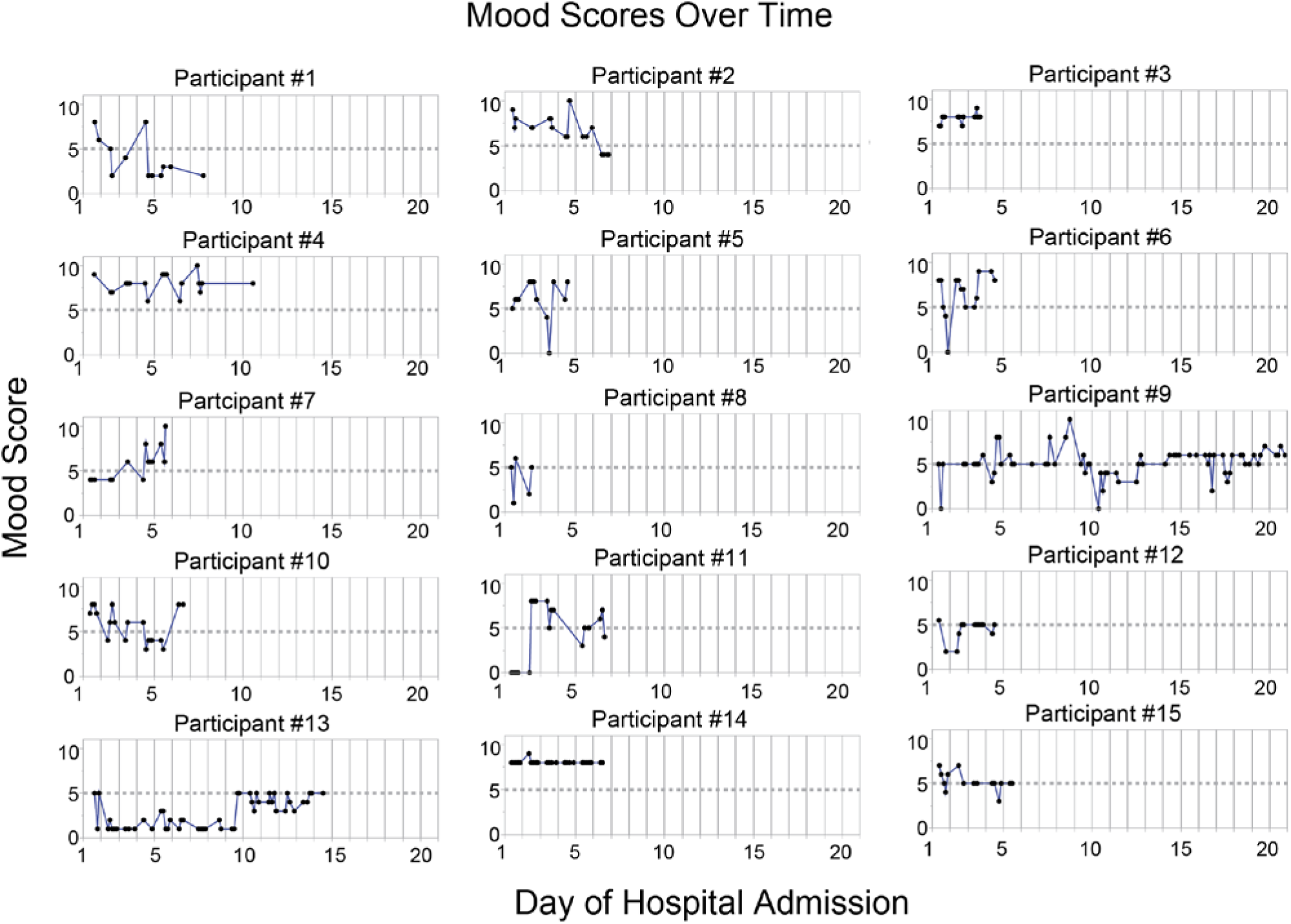
Mood scores over time.

**Figure S2.**
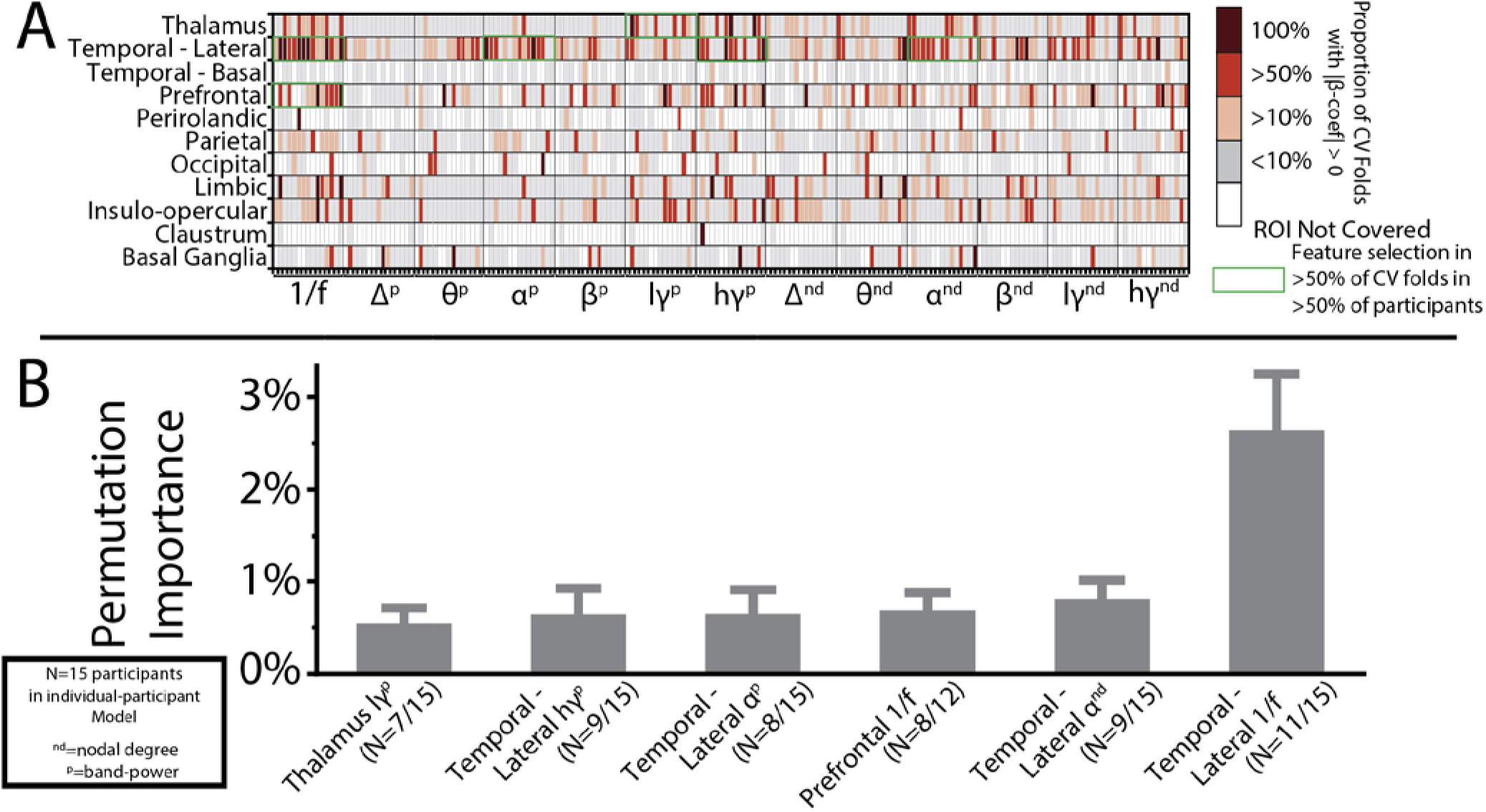
Individual model feature selection (A) and permutation importance (B) when using intracranial encephalography to decode smiles vs neutral expressions. (A) Aperiodic activity, high-gamma (hγ) power, and delta (δ) power in the lateral temporal cortex, along with lγ power in the thalamus were consistently selected in >50% of CV folds and in >50% of participants during individual modeling (green box). (B) Aperiodic activity in the lateral temporal cortex had the sole highest permutation importance (i.e., shuffling feature values associated with each feature, individually and independently, in the optimal model and repeating prediction resulted in at least a 1% drop in accuracy) amongst the consistently selected features.

**Figure S3.**
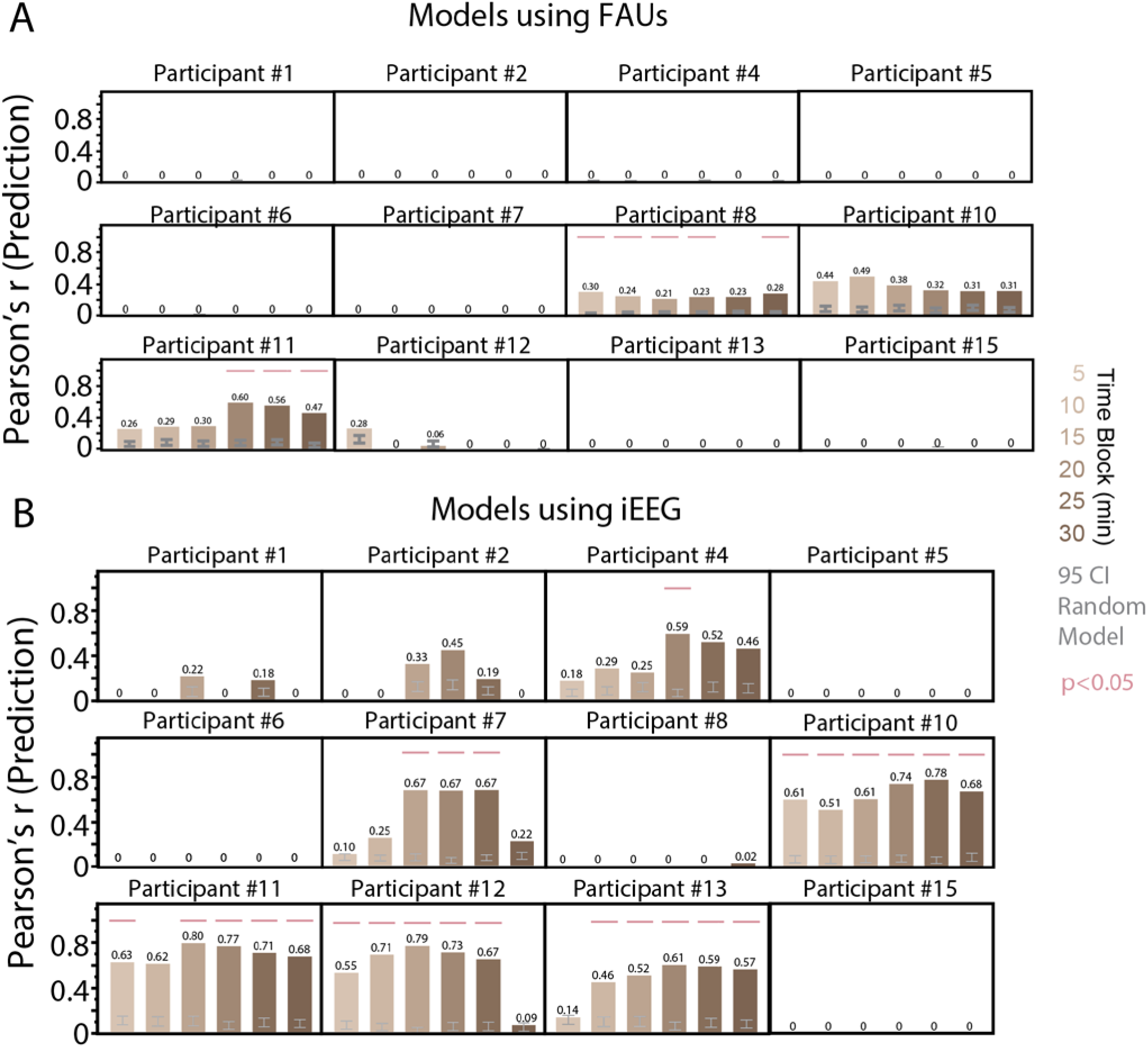
Individual participant model performance in linear regression models using facial action units (FAUs; A) and using intracranial encephalography signals (iEEG) **(B)** to decode mood scores across incremental time blocks surrounding completion of mood survey using Pearson’s r (bar plots), compared to random models (grey error bar). (A) Only 2 of the 12 participants demonstrated consistent decoding using FAUs (significant Pearson’s r in >50% of consecutive time blocks). (B) Only 5 of the 12 participants demonstrated consistent decoding using iEEG.

**Table S1.** Participant Demographics and Summary Distributions of Expression and Mood Data.

| Participant | Age | Sex | BDI | N Neutral | N Smiles | N Mood Scores | Range of Mood Scores | N Unique Mood Scores | log10(N) Unique Permutations of Mood Scores | Smile Exclusion | Mood Exclusion |
| --- | --- | --- | --- | --- | --- | --- | --- | --- | --- | --- | --- |
| #1 | 24 | M | 12 | 404 | 159 | 12 | 6 | 6 | 6 |  |  |
| #2 | 37 | M | 19 | 427 | 131 | 18 | 6 | 6 | 10.2 |  |  |
| #3 | 32 | F | 9 | 307 | 134 | 12 | 2 | 3 | 3.3 |  | Exclude |
| #4 | 36 | M | 21 | 427 | 70 | 18 | 4 | 5 | 8.7 |  |  |
| #5 | 28 | F | 21 | 344 | 39 | 13 | 8 | 5 | 5.6 |  |  |
| #6 | 38 | F | 11 | 518 | 66 | 15 | 9 | 7 | 8.7 |  |  |
| #7 | 28 | F | 39 | 374 | 170 | 14 | 6 | 4 | 5.7 |  |  |
| #8 | 50 | F | 1 | 432 | 103 | 5 | 5 | 4 | 1.8 |  | Exclude |
| #9 | 35 | M | 37 | 375 | 99 | 70 | 10 | 9 | 45.6 |  |  |
| #10 | 32 | M | 18 | 455 | 108 | 19 | 5 | 5 | 10.2 |  |  |
| #11 | 59 | M |  | 396 | 68 | 18 | 8 | 7 | 10.8 |  |  |
| #12 | 27 | M | 15 | 231 | 81 | 13 | 3.5 | 4 | 4.6 |  |  |
| #13 | 23 | M | 18 | 233 | 44 | 50 | 4 | 5 | 29.3 |  |  |
| #14 | 45 | F | 16 | 281 | 51 | 26 | 1 | 3 | 1.4 |  | Exclude |
| #15 | 27 | M |  | 360 | 73 | 15 | 4 | 5 | 6 |  |  |
| #16 | 56 | F | 21 | 532 | 6 | 18 | 3 | 4 | 7.4 | Exclude | Exclude |

